# Motivational Cognitive Maps for Self-Regulated Autonomous Navigation

**DOI:** 10.1101/2025.03.10.642337

**Authors:** Oscar Guerrero Rosado, Adrián F. Amil, Ismael T. Freire, Martin Vinck, Paul F.M.J. Verschure

## Abstract

The mammalian hippocampal formation plays a critical role in efficient and flexible navigation. Hippocampal place cells exhibit spatial tuning, characterized by increased firing rates when an animal occupies specific locations in its environment. However, the mechanisms underlying the encoding of spatial information by hippocampal place cells remain not fully understood. Evidence suggests that spatial preferences are shaped by multimodal sensory inputs. Yet, existing hippocampal models typically rely on a single sensory modality, overlooking the role of interoceptive information in the formation of cognitive maps. In this paper, we introduce the Motivational Hippocampal Autoencoder (MoHA), a biologically inspired model that integrates interoceptive (motivational) and exteroceptive (visual) information to generate motivationally modulated cognitive maps. MoHA captures key hippocampal firing properties across different motivational states and, when embedded in a reinforcement learning agent, generates adaptive internal representations that drive goal-directed foraging behavior. Grounded in the principle of biological autonomy, MoHA enables the agent to dynamically adjust its navigation strategies based on internal drives, ensuring that behavior remains flexible and context-dependent. Our results show the benefits of integrating motivational cognitive maps into artificial agents with a varying set of goals, laying the foundation for self-regulated multi-objective reinforcement learning.

## I. INTRODUCTION

Navigation is a central capability for autonomous agents, both biological and artificial. In robotics and artificial intelligence, navigation algorithms have primarily relied on exteroceptive sensory information, such as visual, LiDAR, or depth data, to construct maps and determine optimal paths in an environment. These approaches leverage techniques such as simultaneous localization and mapping (SLAM), reinforcement learning, and graph-based path planning to enable efficient movement through space. While these methodologies have demonstrated success in a variety of tasks, they remain fundamentally constrained by their reliance on external sensory cues and predefined objectives. The resulting navigation strategies optimize spatial efficiency but lack a key aspect of biological autonomy: the integration of internal states to dynamically modulate behavior [1].

Unlike artificial agents, which typically have explicitly defined and static goals, biological organisms dynamically adjust goal selection based on their physiological state and changing environmental conditions [2]. For instance, an animal’s motivation to seek food or water emerges from its current metabolic needs rather than a predefined objective. Thus, biological navigation is inherently goal-oriented and driven by an organism’s internal needs [3]. This capacity of biological organisms ensures their survival in dynamic environments and highlights an important limitation in stateof-the-art robot navigation: the absence of adaptive goal-selection mechanisms that integrate both external and internal information.

The mammalian hippocampus is widely recognized as a key brain structure involved in spatial cognition. Lesion studies have shown that hippocampal damage impairs performance in spatial tasks [4], and the discovery of spatially tuned cells [5] supports Tolman’s theory of a cognitive map [6]. This cognitive map theory suggests that animals construct internal representations of their environment to guide behavior rather than simply relying on stimulus-response associations. O’Keefe’s seminal work on place cells [7] further substantiated this idea, demonstrating that hippocampal neurons encode specific spatial locations in an environment. Notably, this work also demonstrated that the firing activity of hippocampal neurons is not solely influenced by spatial cues, but that they exhibit rate remapping as a function of features of sensory cues and behavioral task variables [8], [9], and that they can be sensitive to visual and tactile information (See [10] for more recent evidence in the auditory domain).

Beyond spatial encoding, evidence suggests that the hippocampus integrates interoceptive information relevant to an organism’s motivational state. Early studies reported the existence of approach-consummatory cells that differentially fire during eating and drinking behaviors [11], [7]. Kennedy and Shapiro [12] identified motivational place cells in the hippocampus, which alter their activation patterns depending on whether an animal is food- or water-deprived. These findings suggest that hippocampal neurons do not merely encode the organism’s spatial location but also track the organism’s internal state, allowing for adaptive behavior selection based on physiological needs.

Building on this understanding, homeostatic and allostatic principles [13], [14], [15] have explained behavior as a dynamic regulation of the organism’s internal milieu. Models grounded in homeostatic and allostatic principles have successfully captured the hypothalamic regulation of motivation, enabling artificial agents to self-regulate their goals based on their internal state and the dynamic state of the environment [16], [17]. However, these models do not account for how cognitive maps develop and support adaptive navigation. Similarly, models aiming to explain hippocampal function have successfully captured cognitive map features without explaining the influence of motivational states [18], [19], [20]. A promising approach is provided by homeostatic reinforcement learning models. These models ground the reward value of stimuli in the internal state of the agent, facilitating more biologically realistic learning and decision-making processes [19], [21], [22]. Yet, while these approaches contribute valuable insights, they still rely on predefined representations of cognitive maps and do not fully capture the adaptive, motivationally driven nature of hippocampal spatial encoding. Our approach builds upon the idea that the hippocampus compresses sensory information into meaningful representations, a function well captured by biologically-inspired sparse autoencoders [23], [24], [25], [26]. To capture this process, we introduce a motivational hippocampal autoencoder (MoHA) model that integrates interoceptive (motivational) and exteroceptive (visual) information to generate compressed state representations. This architecture enables an artificial agent to encode not only spatial locations but also how these locations relate to its internal needs. To assess whether these internal representations support goal-directed behavior in navigation, we integrate MoHA with Sequential Episodic Control (SEC) [20], an episodic reinforcement learning algorithm designed to enhance decision-making by storing and retrieving structured memory traces. By coupling MoHA’s motivation-sensitive cognitive maps with SEC’s experience-based action selection, we provide a framework for embodied, self-regulated artificial intelligence that draws inspiration from biological autonomy, where past experiences and internal states jointly shape adaptive navigation strategies.

## II. METHODS

For motivational cognitive maps, we adopt the sparse autoencoder approach [26] and expand it to account for motivational information alongside the original visual input (Fig. 1a). In this manner, the objective function of our model minimizes the reconstruction error for both visual and motivational inputs:

**Fig. 1.**
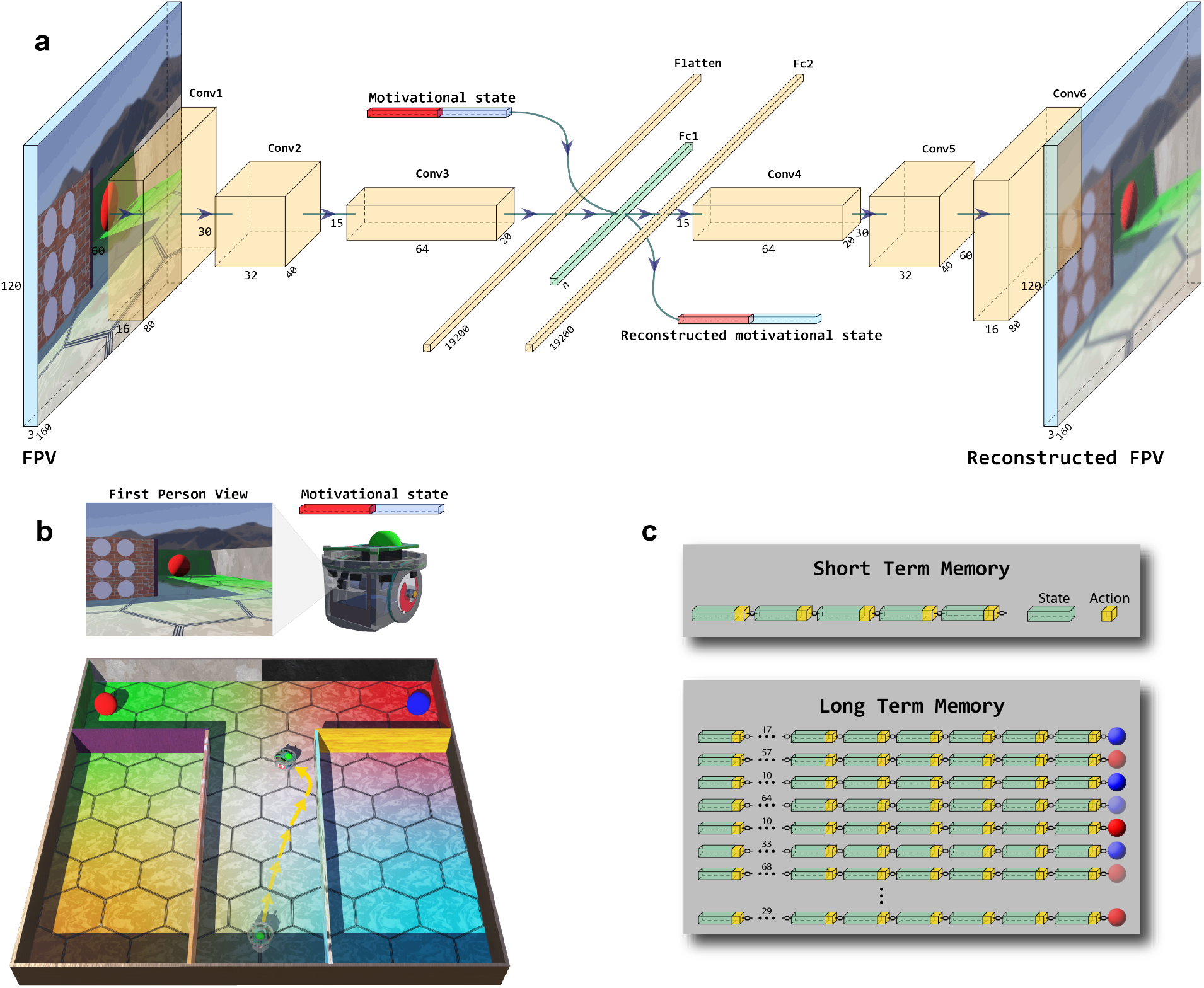
Motivational hippocampal autoencoder (MoHA). a) The MoHA uses the robot’s First Person View (FPV) and its motivational state as input to build a joint compressed representation in its latent vector (green). Based on the latent vector, the decoding side of the network builds reconstructions of the inputs that, in turn, drive error backpropagation. b) Every trial of the spatial task starts with the agent at the beginning of the central alley and with a random dominant motivation. Through actions, the agent navigates the environment, sampling it until it finds a reward. At every step, its visual and motivational experience is passed through the MoHA to generate joint compressed representations of its state. c) These states, together with the actions that led to them, are stored in a short-term memory buffer as couplets. Every new couplet is linked to its predecessor, forming sequences that encode the current behavioral policy. If this plan results in reward gathering, the sequence stored in short-term memory moves to long-term memory together with the obtained reward value. Note that shorter trajectories result in a higher reward, as indicated by the color intensity of the reward stimuli.

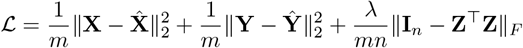

where the first and second terms represent the mean squared error (MSE) between inputs and their reconstructions, **X** denoting visual and **Y** motivational inputs, encouraging the latent vector **Z** to preserve input information. The second term is an orthonormal activity regularization term, whose strength is controlled by *λ*, pushing the Gramian **Z**^⊤^**Z** towards the identity matrix **I**_*n*_. Finally, *m* denotes the batch size and *n* the number of neurons in **Z** (See [26] for a complete description of a regularized sparse autoencoder).

In the navigational task, Sequential Episodic Control (SEC) uses the autoencoder’s latent vectors as the agent’s state to create and optimize efficient behavioral policies. As the agent navigates the environment (Fig. 1b), every state is coupled with the action that led to that state, and a sequential representation of the current behavioral policy is created by linking these state-action couplets. These sequences are maintained in a short-term memory (STM) buffer. Once the agent finds a rewarding stimulus, this sequence is stored in a long-term memory (LTM) system together with the corresponding reward value (Fig. 1c).

Through experience and increased familiarity with the environment, the agent begins encountering states similar to those reward-associated states stored in LTM. As a result, it transitions from random exploration to memory-guided navigation by retrieving actions that proved beneficial in the past. This environmentally mediated synergy between perception and behavior has been shown to systematically bias the agent’s input sampling, which in turn stabilizes behavior [27], [19], [20]. Both STM and LTM systems follow a First-In-First-Out rule once these memory systems reach a parametrically defined storage limit (See [20] for a complete description of Sequential Episodic Control).

Two distinct experiments were conducted to evaluate MoHA’s computational properties, both as an independent model and as an integrated component within a cognitive architecture. First, we train the MoHA with images gathered during random exploration in a Webots [28] simulated Tmaze. Each of these images was randomly associated with one of two possible motivational states. To gain an understanding of the functioning of the MoHA, different model configurations were analyzed. Specifically, we considered the effects of the environment’s visual complexity, embedding size (i.e. the number of units in the latent space), and the motivational vector size. Second, a pre-trained MoHA was embedded in a simulated e-puck mobile robot that used SEC to learn appropriate goal-oriented behavioral policies. This agent started the experiment with empty STM and LTM systems and used spatial and motivational information to improve its navigation to the correct reward site, which was conditional on its current motivational state.

## III. RESULTS

### A. Motivational Cognitive Maps

In order to identify motivationally tuned units in the latent vector of the MoHA we computed a Gain Modulation Score (GMS) for each unit in the latent vector as:

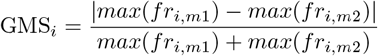

where *fr*_*i*_ is the firing rate of unit *i* and _*m*_ the motivational state for which the firing rate was computed.

The distribution of GMSs across units exhibited a trimodal pattern, suggesting the presence of distinct subpopulations with different modulation properties. Specifically, the histogram revealed peaks at both low and high gain modulation values, with relatively fewer units exhibiting intermediate modulation levels around GMS ≈ 0.5. Notably, a pronounced accumulation of units at GMS ≈ 1 indicated a substantial subset of units that were fully suppressed in one of the two conditions. Fig. 2a shows the distribution of GMSs for a reference model trained in the T-maze v2 (Appx. B) with a latent vector of 400 units and a motivational input size of 10^3^.

**Fig. 2.**
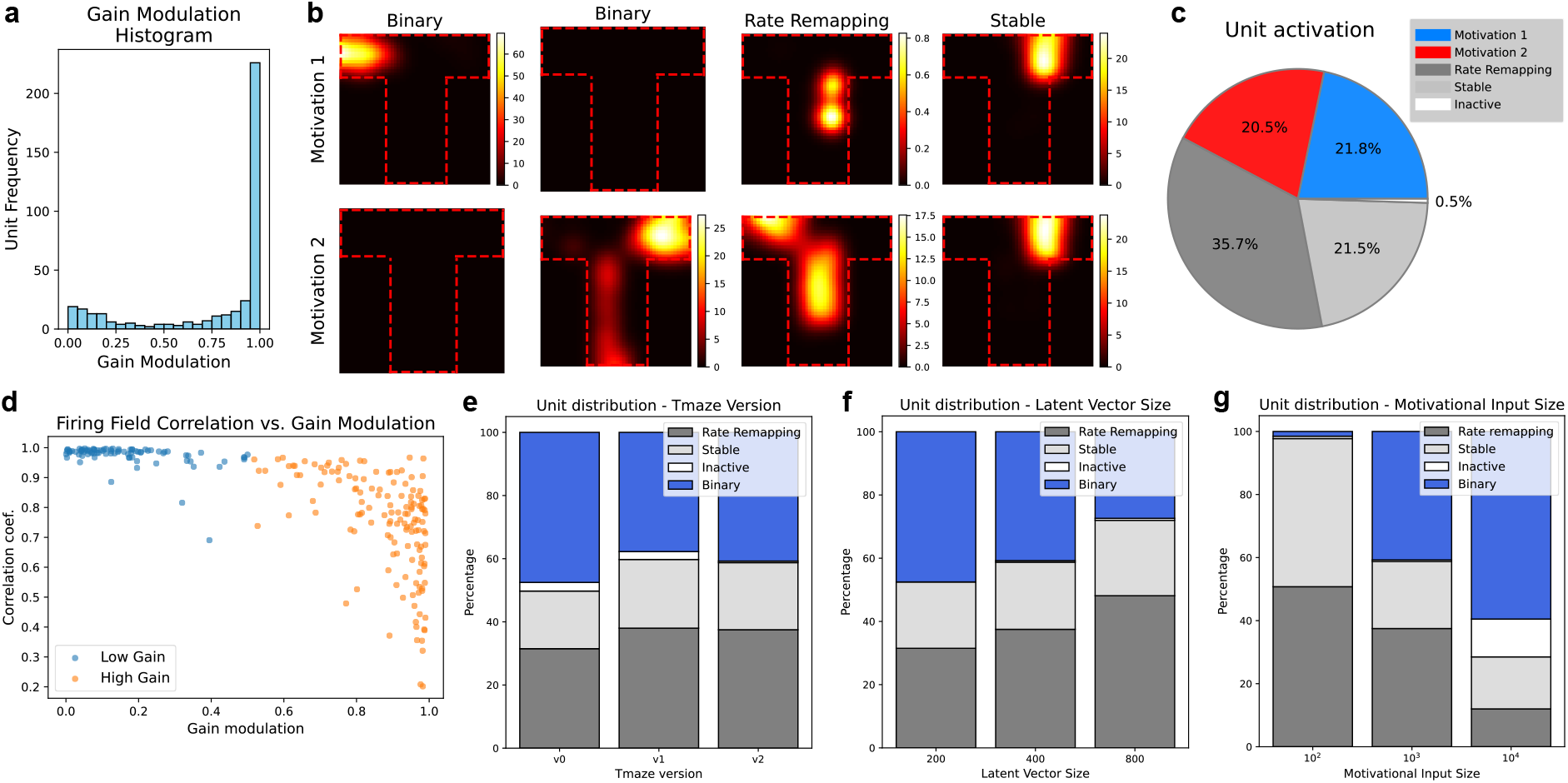
Motivational Cognitive Maps. a) Histogram of Gain Modulation Scores (GMSs) for a reference model trained in T-maze v2 with 400 units in its latent vector and motivational input size of 10^3^. b) Rate maps of example units during motivation 1 and 2 separately. Notice that binary units display activation for only one of the two motivational states. Rate remapping units may get active in a similar location but with a distinct firing rate. Firing fields of stable units remain unaffected by motivational states. c) Unit class distribution for the reference model. Binary units were sub-classified based on their preferential activation to one of the two motivational states. d) Firing field correlation is related to Gain Modulation Scores. As units display lower gain modulation, they become more correlated across motivational states. e) Unit distribution comparison across T-maze versions (Appx. B). f) Unit distribution comparison across latent vector size. g) Unit distribution comparison across motivational input size.

Based on the observed distribution, we categorized units into three distinct groups according to their modulation characteristics (Fig. 2b):

- **Binary units** with GMS *>* 0.99, i.e. active in one motivational context while almost completely inactive in the other.
- **Rate remapping units** with 0.5 ≤ GMS ≤ 0.99, showing intermediate levels of gain modulation and re-flecting changes in firing rate between contexts without complete suppression or activation.
- **Stable units:** with GMS *<* 0.5, exhibiting minimal modulation between contexts, such that their firing properties remain relatively stable.

This classification method differs from that used by Kennedy and Shapiro [12], which used firing-field correlations to quantify cell remapping. However, the unit class distribution of our reference model still resembles that found in physiological data (Fig. 2c) and, when comparing gain modulation and field correlation, low gain-modulated (i.e. stable) units showed high correlation values (Fig. 2d). Thus, when encoding states under two different motivational contexts, our classification approach not only captures global remapping, in which place fields shift to other locations, but also rate remapping, where firing rates are gain-modulated. Indeed, rate remapping has been reported under different conditions, such as the discrimination of different sensory cues [8] and when analyzing social place cells [9].

To determine whether visual information affects unit class distribution in the MoHA, we trained the model under three spatially identical yet visually distinct T-maze configurations. Our results do not show an apparent alteration of the unit class distribution driven solely by visual differences in the environment (Fig. 2e) (Appx. B). However, we did find that the unit class distribution was sensitive to the architectural features of the model. The percentage of binary units tended to decrease as we increased the size of the latent vector (Fig. 2f); an effect accompanied by an increase in the proportion of rate-remapping units. The opposite effect occurred when increasing the size of the motivational input (Fig. 2g).

### B. A motivationally-guided SEC agent

To elucidate whether MoHA-generated motivational cognitive maps are adequate to guide spatial learning, we integrate them with Sequential Episodic Control (SEC). A simulated mobile robot (i.e., e-puck) was equipped with this architecture to learn goal-directed navigation strategies in a spatial decision task, where motivational information played a key role in selecting the correct target.

To compare performance, a control robot was equipped with an architecture comprising a hippocampal autoencoder trained without motivational inputs. The experiment was divided into two parts: First, both control autoencoder and MoHA were pre-trained with images gathered during random exploration of the environment. As in the previous experiment, the MoHA paired each of these images with a random motivation. Second, SEC enabled learning in the T-maze foraging task. Here, both motivational and control agents shared the same STM and LTM of size 50 and 200, respectively.

During the pretraining of the motivational autoencoder, visual reconstruction error remained lower for the control model, which is an expected result since the control autoencoder did not dedicate resources for the encoding of motivational information as in the case of the MoHA (Fig 3a). Although differences in the reconstruction error were visually appreciable, both the control and motivational autoencoder models reconstructed the input images successfully (Fig 3b). Moreover, when analyzing the unit’s spatial information [29] (Appx. A), the control autoencoder did not differ from the motivational autoencoder during motivation 1 or 2 (Fig 3c).

**Fig. 3.**
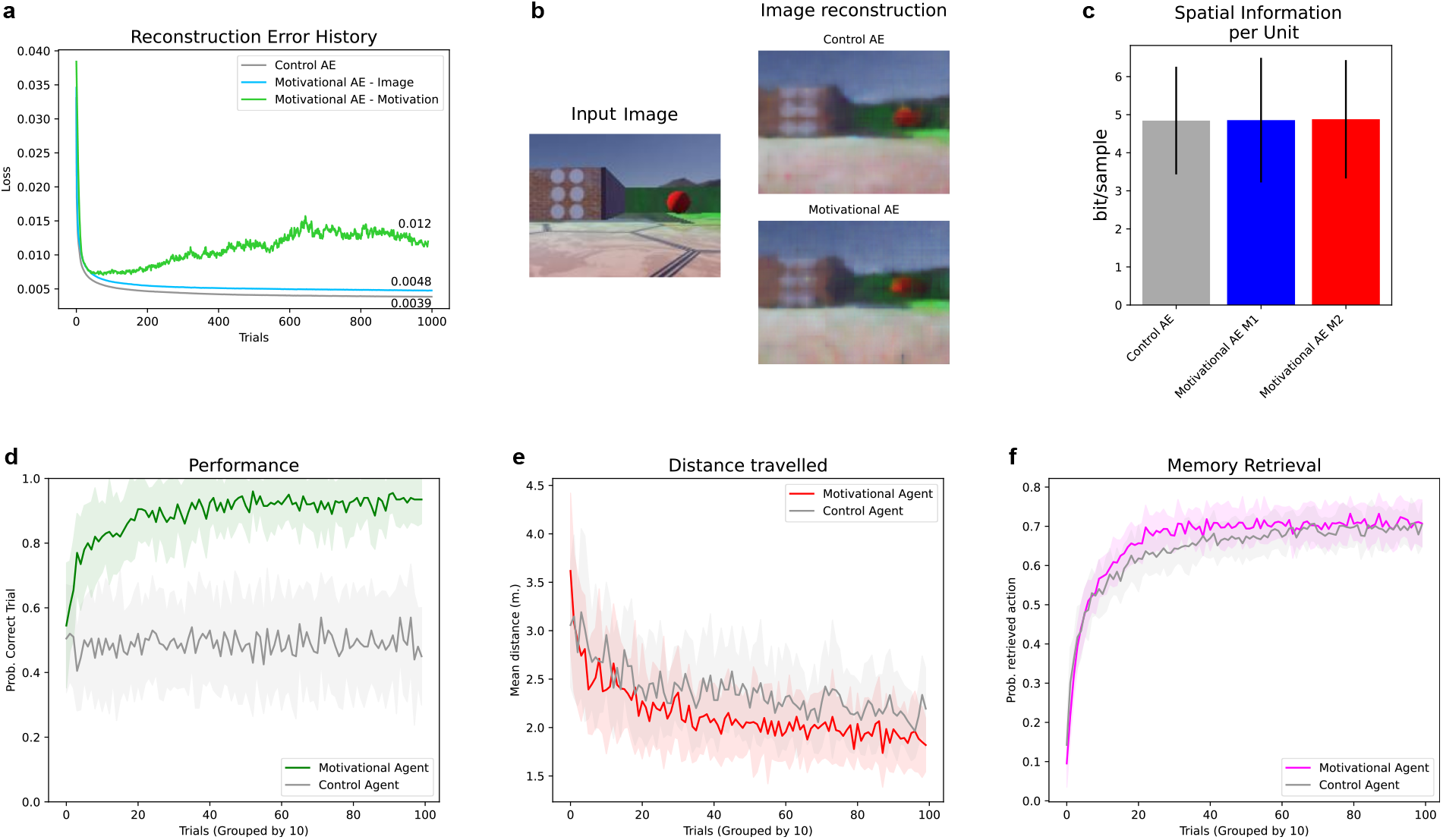
Performance of a motivationally-guided artificial agent in the Tmaze foraging task. a) Reconstruction error history for both control and MoHAs (Motivational Hippocampal Autoencoders). b) Image reconstruction example for both control and MoHA. c) Spatial information comparison. Splitted into motivations 1 and 2 for the MoHA. d) Agents’ performance as the probability of gathering the correct reward (given the trial’s dominant motivation). e) Trajectory optimization across trials. f) Memory retrieval measured as the probability of performing a memory-guided action over a random one. Plots d, e, and f inform about mean measures gathered during 20 experiments for both motivational and control agents.

In the spatial decision-making task, where both agents started every trial with a random dominant motivation, only the motivationally-guided agent successfully learned to navigate to the correct reward. After approximately a hundred trials, the agent successfully navigated to the correct reward more than 80 percent of the time, and its probability of choosing the correct reward kept improving (Fig. 3d). In contrast, the control agent showed poor performance, with a probability of choosing the correct reward around chance levels during the entire experiment. Despite this difference in performance, both motivational and control agents learned to optimize their trajectories toward rewards (Fig. 3e) and increased the rate of memory-guided actions performed along the trial (Fig. 3f). These results demonstrate that although the control agent successfully learned to navigate toward reward sites, only the motivational agent could adapt its selected target location based on the currently dominant motivational context.

## IV. DISCUSSION

In the present work, we developed a motivational hippocampal autoencoder (MoHA) that jointly encoded motivational and spatial states. This joint encoding led to the development of motivational-context-dependent place fields, exhibiting rate remapping.

Hippocampal rate remapping has been previously described based on changing sensory cues or the presence of another agent in the same environment [8], [9]. This phenomenon highlights the key distinction between global remapping, in which place fields shift to other locations, and rate remapping, in which firing rates are gain-modulated [8].

Our results strongly indicate that MoHA exhibits rate remapping, as its unit distributions showed gradual gain modulation rather than full shifts in firing locations when encoding different motivational states. A rate-based classification approach allowed us to identify binary, stable, and rate remapping units in a distribution similar to that found in physiology by Kennedy and Shapiro [12]. Nonetheless, their classification approach was based on place-field correlations, an approach that cannot distinguish rate remapping from global remapping. Further experiments are needed to determine whether MoHA could also support global remapping. A potential future direction is to test whether the integration of a FiLM layer [30] within the MoHA drives global remapping while conserving spatially constrained firing fields. In addition to exploring MoHA’s neural representations, we have tested its suitability in motivationally guided foraging tasks, where different rewards must be acquired based on the internal state of the agent. Our integration of MoHA with Sequential Episodic Control (SEC) [20] leveraged MoHA’s state representations to generate optimal behavioral policies. This resulted in efficient performance in the T-maze foraging task, demonstrating that motivational cognitive maps are necessary for an agent to learn reward-directed strategies.

The interoceptive modulation of hippocampal activity remains an open question, with potential mechanisms involving the hippocampus’s unique connectivity and receptor expression. The hippocampus interfaces with the lateral ventricle, providing access to internal markers such as hormones, metabolites, and temperature fluctuations. Additionally, hippocampal neurons express a diverse array of receptors, far more than the average neuron, allowing them to integrate multimodal sensory and interoceptive signals [31]. This extensive receptor profile enables the hippocampus to link bodily states to cognitive processes. Understanding how these mechanisms contribute to the formation of motivational cognitive maps is crucial for advancing biologically inspired artificial intelligence.

Beyond its neuroscientific relevance, our findings have direct implications for robotic autonomy. Most state-of-theart approaches in robot navigation rely on externally defined goals and operate under static reward structures, limiting their ability to dynamically adapt to changing internal states. By contrast, MoHA enables an agent to generate motivational cognitive maps, where goal selection is contingent on its motivational state. This represents a step toward biologically inspired robotic autonomy, allowing for more adaptive and goal-directed decision-making.

MoHA potentially addresses two core challenges in robotics: First, it enables robots to prioritize competing goals and separate strategy representations based on their internal states, partially solving task management and action orchestration problems. Second, by continuously updating cognitive maps based on both sensory and motivational inputs, MoHA provides a foundation for lifelong reinforcement learning in dynamic environments, where an agent can adapt its decision-making over time. Future work should be dedicated to exploring whether MoHA could encode a larger set of motivational states, opening the door to biologically constrained multi-objective reinforcement learning. The scalability of MoHA to larger real-world scenarios also deserves future investigation.

## ACKNOWLEDGMENT

This study was funded by Counterfactual Assessment and Valuation for Awareness Architecture—CAVAA (European Commission, EIC 101071178)

## APPENDIX

## A. Spatial information

Spatial information is computed as:

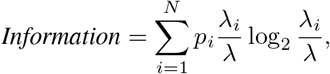

where *i* = 1, …, *N* denotes nonoverlapping spatial bins dividing the environment, *p*_*i*_ is the occupancy probability of bin *i* given the agent’s random sampling distribution, *λ*_*i*_ is the mean unit activation for bin *i*, and *λ* is the overall mean activation of the unit in the environment.

### B. T-maze versions

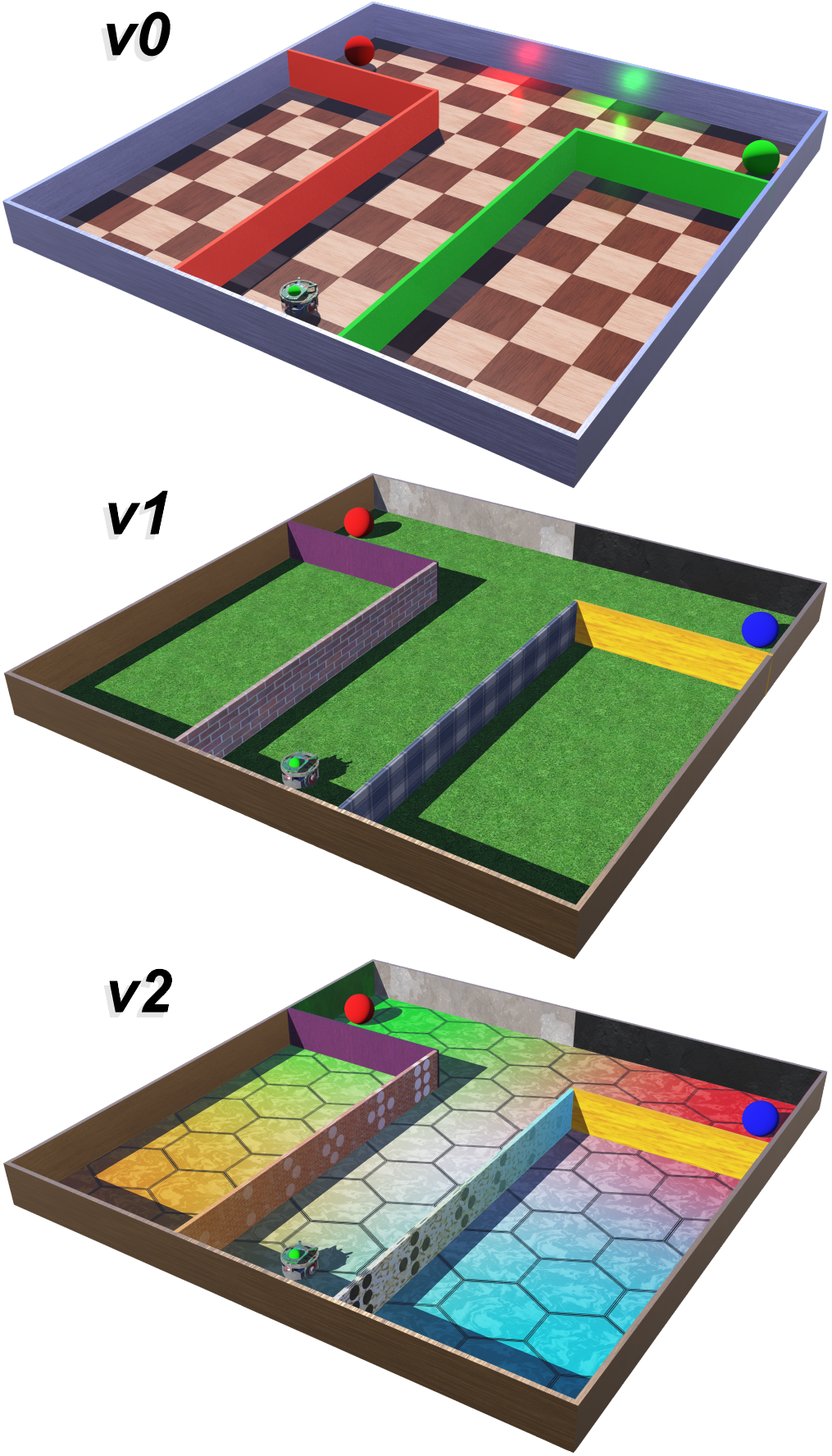

## REFERENCES

[1] Matteo Mossio and Alvaro Moreno. Biological autonomy: A philosophical and theoretical enquiry. history, philosophy, and theory of life sciences series, 2015.

[2] Clark L Hull. Principles of behavior, 1945.

[3] Margaret Wilson. Six views of embodied cognition. Psychonomic bulletin & review, 9:625–636, 2002.

[4] Robert J Douglas. The hippocampus and behavior. Psychological bulletin, 67(6):416, 1967.

[5] John O’Keefe and Jonathan Dostrovsky. The hippocampus as a spatial map: preliminary evidence from unit activity in the freely-moving rat. Brain research, 1971.

[6] Edward C Tolman. Cognitive maps in rats and men. Psychological review, 55(4):189, 1948.

[7] John O’Keefe. Place units in the hippocampus of the freely moving rat. Experimental neurology, 51(1):78–109, 1976.

[8] Stefan Leutgeb, Jill K Leutgeb, Carol A Barnes, Edvard I Moser, Bruce L McNaughton, and May-Britt Moser. Independent codes for spatial and episodic memory in hippocampal neuronal ensembles. Science, 309(5734):619–623, 2005.

[9] Jeroen J Bos, Martin Vinck, Pietro Marchesi, Amos Keestra, Laura A van Mourik-Donga, Jadin C Jackson, Paul FMJ Verschure, and Cyriel MA Pennartz. Multiplexing of information about self and others in hippocampal ensembles. Cell Reports, 29(12):3859–3871, 2019.

[10] Dmitriy Aronov, Rhino Nevers, and David W Tank. Mapping of a non-spatial dimension by the hippocampal–entorhinal circuit. Nature, 543(7647):719–722, 2017.

[11] James B Ranck Jr. Studies on single neurons in dorsal hippocampal formation and septum in unrestrained rats: Part i. behavioral correlates and firing repertoires. Experimental neurology, 41(2):462–531, 1973.

[12] Pamela J Kennedy and Matthew L Shapiro. Motivational states activate distinct hippocampal representations to guide goal-directed behaviors. Proceedings of the National Academy of Sciences, 106(26):10805–10810, 2009.

[13] Claude Bernard. Introduction á l’étude de la médecine expérimentale par m. Claude Bernard. Baillière, 1865.

[14] Walter Bradford Cannon. The wisdom of the body. 1939.

[15] Peter Sterling. Allostasis: a new paradigm to explain arousal pathology. Handbook of life stress, cognition and health, 1988.

[16] Marti Sanchez-Fibla, Ulysses Bernardet, Erez Wasserman, Tatiana Pelc, Matti Mintz, Jadin C Jackson, Carien Lansink, Cyriel Pennartz, and Paul FMJ Verschure. Allostatic control for robot behavior regulation: a comparative rodent-robot study. Advances in Complex Systems, 13(03):377–403, 2010.

[17] Oscar Guerrero Rosado, Adrian F Amil, Ismael T Freire, and Paul FMJ Verschure. Drive competition underlies effective allostatic orchestration. Frontiers in Robotics and AI, 9:1052998, 2022.

[18] Marti Sanchez Fibla, Ulysses Bernardet, and Paul FMJ Verschure. Allostatic control for robot behaviour regulation: An extension to path planning. In 2010 IEEE/RSJ International Conference on Intelligent Robots and Systems, pages 1935–1942. IEEE, 2010.

[19] Giovanni Maffei, Diogo Santos-Pata, Encarni Marcos, Marti Sánchez-Fibla, and Paul FMJ Verschure. An embodied biologically constrained model of foraging: from classical and operant conditioning to adaptive real-world behavior in dac-x. Neural Networks, 72:88–108, 2015.

[20] Ismael T Freire, Adrián F Amil, and Paul FMJ Verschure. Sequential memory improves sample and memory efficiency in episodic control. Nature Machine Intelligence, pages 1–13, 2024.

[21] Mehdi Keramati and Boris Gutkin. Homeostatic reinforcement learning for integrating reward collection and physiological stability. Elife, 3:e04811, 2014.

[22] Hugo Laurençon, Charbel-Raphaël Ségerie, Johann Lussange, and Boris S Gutkin. Continuous homeostatic reinforcement learning for self-regulated autonomous agents. arXiv preprint 2109.06580, 2021.

[23] Marcus K Benna and Stefano Fusi. Place cells may simply be memory cells: Memory compression leads to spatial tuning and history dependence. Proceedings of the National Academy of Sciences, 118(51):e2018422118, 2021.

[24] Diogo Santos-Pata, Adrián F Amil, Ivan Georgiev Raikov, César Rennó-Costa, Anna Mura, Ivan Soltesz, and Paul FMJ Verschure. Entorhinal mismatch: A model of self-supervised learning in the hippocampus. Iscience, 24(4), 2021.

[25] Diogo Santos-Pata, Adrián F Amil, Ivan Georgiev Raikov, César Rennó-Costa, Anna Mura, Ivan Soltesz, and Paul FMJ Verschure. Epistemic autonomy: self-supervised learning in the mammalian hippocampus. Trends in cognitive sciences, 25(7):582–595, 2021.

[26] Adrian F Amil, Ismael T Freire, and Paul FMJ Verschure. Discretization of continuous input spaces in the hippocampal autoencoder. arXiv preprint 2405.14600, 2024.

[27] Paul FMJ Verschure, Thomas Voegtlin, and Rodney J Douglas. Environmentally mediated synergy between perception and behaviour in mobile robots. Nature, 425(6958):620–624, 2003.

[28] Webots. http://www.cyberbotics.com. Open-source Mobile Robot Simulation Software.

[29] William Skaggs, Bruce Mcnaughton, and Katalin Gothard. An information-theoretic approach to deciphering the hippocampal code. Advances in neural information processing systems, 5, 1992.

[30] Ethan Perez, Florian Strub, Harm De Vries, Vincent Dumoulin, and Aaron Courville. Film: Visual reasoning with a general conditioning layer. In Proceedings of the AAAI conference on artificial intelligence, volume 32, 2018.

[31] György Buzsáki and David Tingley. Cognition from the body-brain partnership: exaptation of memory. Annual review of neuroscience, 46(1):191–210, 2023.

